# Inter- and intra-individual differences in fluid reasoning show distinct cortical responses

**DOI:** 10.1101/039412

**Authors:** Rogier A. Kievit, H. Steven Scholte, Lourens J. Waldorp, Denny Borsboom

## Abstract

Fluid intelligence is a general cognitive ability associated with problem solving in the absence of task-specific knowledge. Neuroscientific studies of fluid intelligence have studied both fluid intelligence tasks of varying difficulty and individual differences in fluid intelligence ability, but have failed to appropriately distinguish the two dimensions. Here we use task-based fMRI (N=34) to show that *within* and *between* subject dimensions show both partial overlap and widespread differences. Individuals with higher ability showed widespread increased activity including bilateral frontoparietal systems, whereas more difficult items were associated with more focal activity increases in middle frontal gyri, frontal poles and superior frontal poles. Finally, we show that when difficulty is equated across individuals, those with higher ability tend to show more fronto-parietal activity, whereas low fluid intelligence individuals tend to show greater activity in higher visual areas. The fMRI and behavioural data for our paper are freely available in online repositories.

## Introduction

Fluid intelligence is the ability to think logically and solve novel problems in the absence of task-specific knowledge (Horn and Cattell, 1966). It is a central component of psychometric theories of intelligence (Carpenter et al., 1990; Carroll, 1993; Engle et al., 1999) and closely related to core cognitive abilities including working memory (Engle et al., 1999), processing speed (Fry and Hale, 1996), attention (Engle, 2002), general intelligence (Blair, 2006) and executive functions (Salthouse et al., 2003). Individuals with higher fluid reasoning ability generally have better psychosocial outcomes (Strenze, 2007; Deary, 2012), lower instances of psychopathology (Gale et al., 2010) and lower morbidity and mortality (Deary et al., 2011). Moreover, fluid intelligence often declines rapidly in old age (Kievit et al., 2014; Aichele et al., 2015) with adverse consequences for the ability to live and function independently (Salthouse et al., 2003; Tucker-Drob, 2011).

Cognitive neuroscience has contributed a variety of insights into the neural processes and properties associated with fluid reasoning, including individual differences in (fluid) intelligence (Choi et al., 2008; Deary et al., 2010; Cole et al., 2012; Kievit et al., 2012, 2014; Ritchie et al., 2015), neural responses during fluid reasoning tasks of varying complexity (e.g. Duncan, 2000; Gray et al., 2003; Lee et al., 2006) and the adverse effects of lesions (Duncan et al., 1995; Roca et al., 2010). Together, these findings converge on a distributed parietal and frontal network associated with fluid reasoning (Kane and Engle, 2002; Jung and Haier, 2007; Fedorenko et al., 2013).

However, studies of fluid intelligence often implicitly conflate two sources of variation: Differences *between* subjects (i.e., differences in ability) and differences *within* subjects (i.e., differences in neural responses under varying task difficulty) (see also (Cronbach, 1957; Chabris, 2007). For instance, the Parieto-Frontal integration model (Jung and Haier, 2007) is a *process model* of reasoning behaviour (p. 138). That is, it claims to describe the processes that happen *within* a subject during complex reasoning. However, it is largely based on neuroimaging studies concerning differences *between* individuals. This is problematic, as it is well known that these dimensions can, and do, behave independently (Hamaker et al., 2005; Penke et al., 2011; Kievit et al., 2013). This leaves a fundamental ambiguity in what is meant, exactly, by the ‘neural substrate’ of fluid reasoning (e.g. Prabhakaran et al., 1997). Does this term refer to the question which neural systems are *differentially recruited* depending on the complexity of the task, or to which neural systems are *differentially active* between people of differing fluid reasoning ability? By not addressing the two dimensions of difficulty and ability simultaneously, studies that focus on either dimension implicitly treat the other dimension of variation as a source of noise, affecting the findings to an unknown degree. Understanding this distinction in detail is crucial to our understanding of both the process of fluid reasoning and individual differences in fluid reasoning ability.

In the present paper, we use Item Response Theory (IRT, Embretson and Reise, 2013) to decompose neural responses during a fluid reasoning task into an inter-individual dimension and an intra-individual dimension. An IRT model combines item difficulty estimates (intra-individual parameters) with ability estimates for each subject (an inter-individual parameter). Using this model, we can separate neural systems that underlie individual differences those that reflect differences in increasing task difficulty. We hypothesize that the neural networks that are differentially active within people with differing ability are not the same as neural networks that are differentially active within people across tasks of varying difficulty. Crucially, by taking into account both dimensions we can compare individual differences in a novel manner: By selecting a differing subset of items for each individual tailored to their ability level, we can compare individual differences in terms of neural activity patterns whilst keeping intra-individual differences in subjective difficulty constant. Doing so sheds new light on the controversial notion of neural efficiency, and illustrates the power of simultaneously modelling inter-and intra-individual differences in a GLM framework.

## Results

### Beha vioural results

To decompose the differential contributions of difficulty and ability in neural response, we fit a Rasch model to the response patterns of a set of Raven's Matrices (see Figure 1 for an example, see materials and methods for more detail).

**Figure 1:**
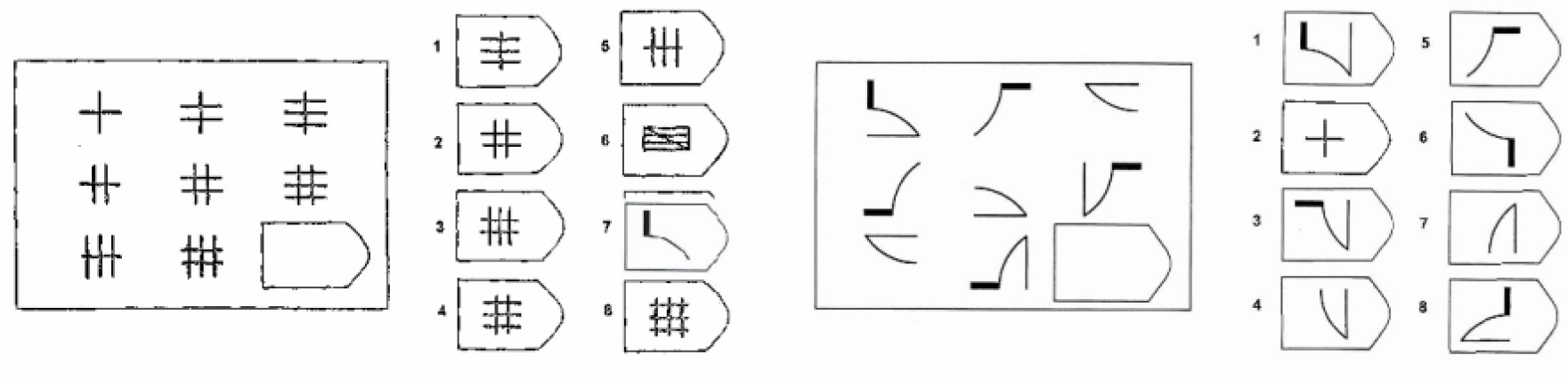
Stylized example an easy (left) and hard (right) Raven's Matrix fluid reasoning task.

A Rasch model is one from a family of Item Response Theory models (IRT; Hambleton et al., 1991; Embretson and Reise, 2013). Models such as the Rasch model can decompose two dimensions simultaneously, better deal with measurement error, explicitly test the assumption that only one ability is being tested, and (in certain cases) a properly specified measurement model can increase statistical power (e.g. Sluis et al., 2010). IRT models remain relatively rarely used in neuroimaging (but see Thomas et al., 2013). In the Rasch model, the difficulty of items is related to the ability of participants by means of a logistic function. The probability that person *j* with ability 9 makes item *i* with difficulty β correctly can be described by equation (1).

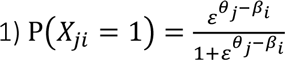

Variants of Rasch models are widely used in fields such as educational testing (Bond and Fox, 2006) and more specific skills such as chess ability (van der Maas and Wagenmakers, 2005). In the Rasch model we model *M* dichotomously scored items (1=correct, 0=incorrect) for *N* persons. Each item has a difficulty parameter β, and each person has an ability parameter θ. We fit a Rasch model in R (Team, 2014) using packages ltm (Rizopoulos, 2006) and eRm (Mair and Hatzinger, 2007). We considered both null-responses (no response within the 30 second time limit) and incorrect responses as incorrect, giving each participant a potential range of 0 to 72 correct. The 34 participants made an average of 39.6 items correct (range: min=19, max=53, SD=8.8). The mean reaction time across individuals was 15.90s, SD= 2.39s, with an item level RT ranging from 1.2 s to a maximum of 29.99 s. To best estimate the ability parameter (θ) of each participant, we fixed the difficulty parameters (β) of the 72 items based on the Ravens standardization sample (Raven, Court, & Raven, 1996). The difficulty parameters of the items ranged from -3.59 to 4.8, capturing a wide range of difficulties. The Andersen Likelihood-Ratio test (Andersen, 1973) indicated that the response pattern fit the Rasch model adequately: χ^2^(46, N=34)= 38.739, p=.767. Figure 2 (top) shows the Item Response Curves of the 72 items and a histogram of the distribution of the ability scores (θ) of the participants (bottom).

To ensure that our sample of participants performed the test accurately, we next fit the model without the item-level constraints, to examine whether the difficulties estimated in our sample matched the difficulties based on the standardization sample. Despite a relatively small sample size, the betas showed a high degree of convergence with published standards (*r* (70) =.85, *p*<.0001). Further analyses showed that more difficult items (with higher betas) were associated with slower response times (Spearman's *r* of reaction times with correct response: *r* (2446) =.55, *p*<.0001), were less likely to be made correctly (Spearman's *r* difficulty with correct response= r (2446) =-.59, *p*<.0001) and were more likely to be null-responses (Spearman's r difficulty with null response = r (2446) =.27, *p*<.0001)

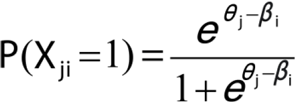

**Figure 2:**
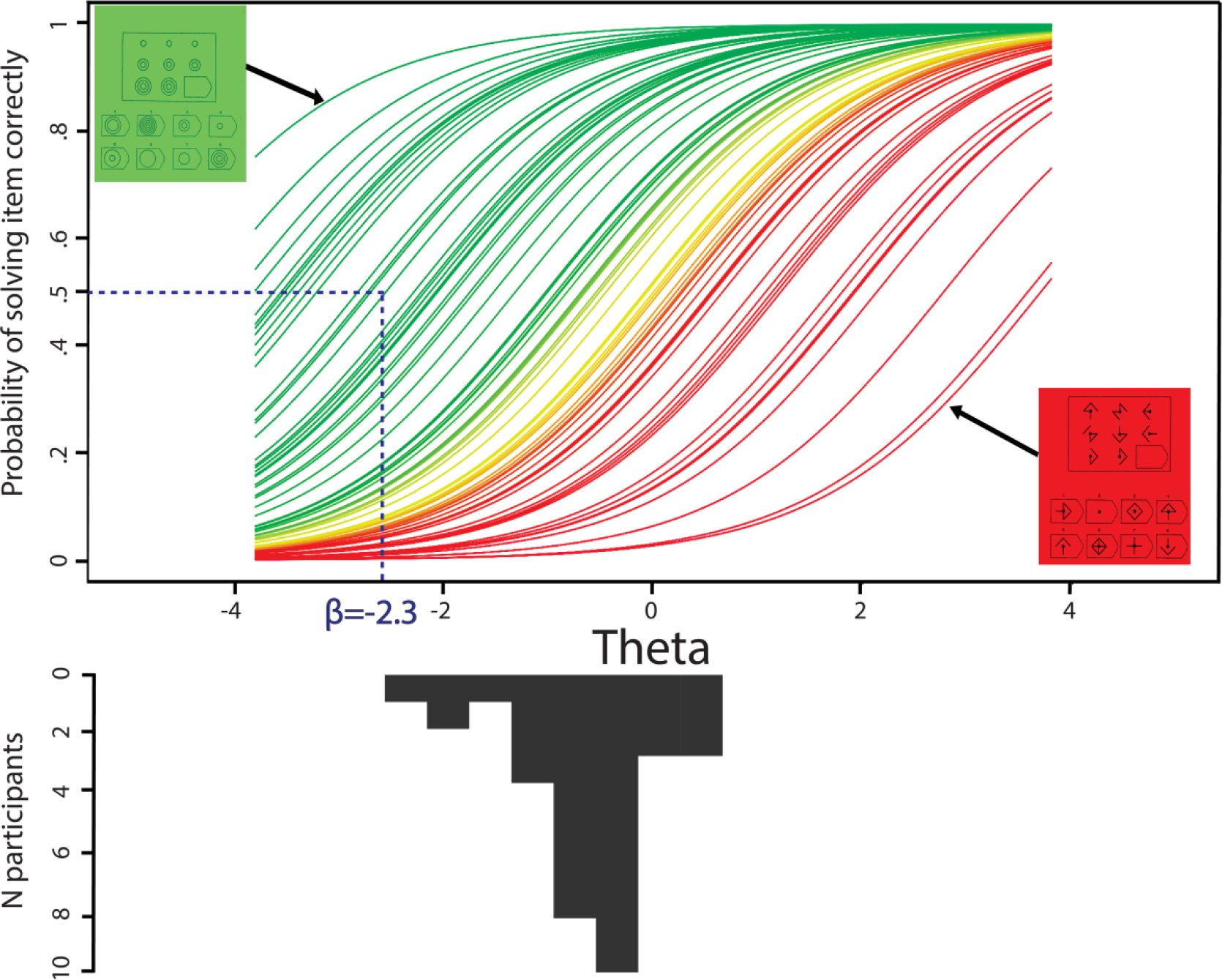
Item-characteristic curves for all 72 items (top). The 72 Raven's matrices items represented as ranging from easy (green/left) to hard (red/right). Ability is modelled such that person parameter theta corresponds to the probability of person j making item i correctly. The difficulty of an item (beta) can be read off by looking up the position on the X-axis that corresponds to a probability of .5 of making that item correctly (example shown in blue).

Together these behavioural analyses suggest that the behavioural manipulation of fluid intelligence was successful: Participants took longer to respond to more difficult items, were less likely to respond correctly and were more likely to fail to respond within the time limit. Moreover, the pattern of responses was well-described by a Rasch model. For all further neuroimaging analyses we use the estimates of difficulty (β, based on the standardization sample) and ability (θ) to study the neural systems underlying differences in item difficulty and ability.

### Differences as a function of item difficulty

First, we examined which regions showed more activity for more difficult items when including all individuals and all difficulty levels. To do so, we take the beta estimates of the difficulty of the 72 items as shown in Figure 2, and used them to predict differential brain activity for each individual, controlling for individual differences in mean activity, with a FLAME random effects analysis. We include both correct and incorrect items, as the cognitive processes that ultimately lead to incorrect answers are as much part of fluid reasoning as the cognitive processes that lead to correct responses. Figure 3 shows five clusters that show increased activity as a function of increasing difficulty. These clusters include the bilateral precunei, the bilateral superior parietal cortices, the superior frontal gyrus (L) and the precuneus bordering on the posterior cingulate. A large parietal cluster was formed by the right and left angular gyri in the parietal lobes (Brodmann areas 39), in line with work showing that activity in as varying parametrically in activity with an increase in complexity (Kroger et al., 2002). Together, this activity pattern is broadly in line with a broad, parieto-frontal network often associated with complex tasks (Jung and Haier, 2007; Duncan, 2010).

**Figure 3/Table 1:**
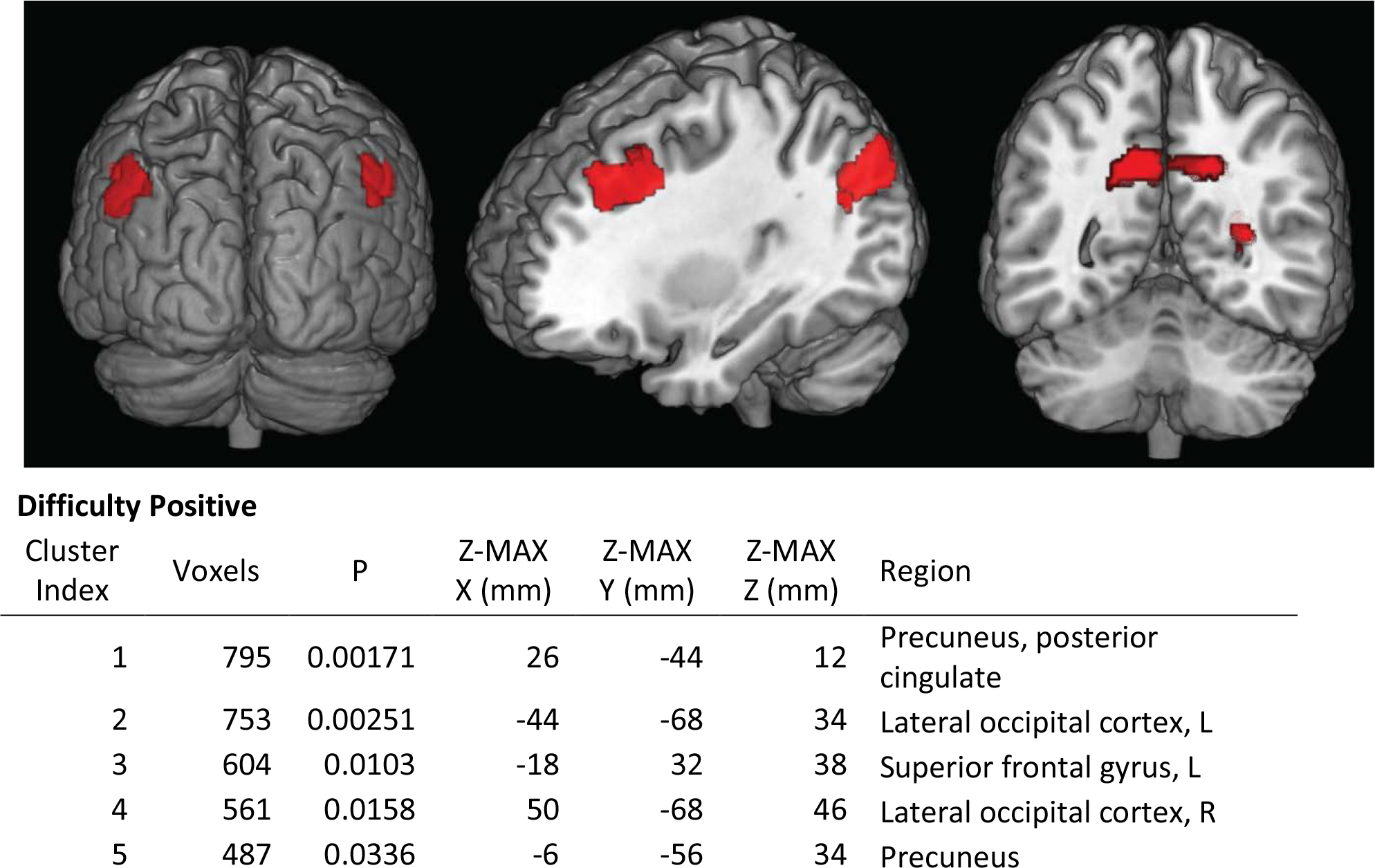
Spatial activity maps associated with an increase in item-difficulty (top), with peak activations of each of the five clusters shown below. No regions showed less activity for more difficult items.

### Individual differences in neural networks during fluid reasoning

Second, we examined individual differences in neural responses during the fluid reasoning task. The ability estimates theta, one for each participant (shown in Figure 2, bottom), were entered into a FLAME random effects analysis to account for individual differences in ability. The results in Figure 4 show that individuals with higher estimated ability showed greater activity in a widespread, bilateral set of regions. The regions of greater activity include bilateral inferior and superior parietal cortices, a large portion of the paracingulate gyrus, bilateral (but greater on the left) Brodmann areas 10, and bilateral middle frontal gyri. These regions have been associated with a wide variety of executive functioning tasks. The large parietal network, consisting of Brodmann areas 7, 39 and 40 (inferior and superior parts of the parietal cortex) have been reported in a variety of imaging studies associated with individual differences in intelligence (e.g. Lee et al., 2006; Choi et al., 2008). The regions of greater activity in individuals with higher ability show overlap with what is known as the Multiple Demand System (Duncan, 2010; Fedorenko et al., 2013). The multiple demand system is a distributed set of regions throughout the cortex known to be differentially active in a wide range of complex tasks such as working memory, interference monitoring, and mathematical problem solving (Fedorenko et al., 2013) both in humans and in single cell recordings in non-human primates (Kusunoki et al., 2009). Lesions to regions within the MDS lead to disproportionate problems in tasks of executive functioning compared to other cognitive abilities (Duncan et al., 1995; Woolgar et al., 2010). Conversely, a set of three regions (shown in Table 2 and Figure 4, yellow) showed more activity in people with lower ability, most notably in bilateral lateral occipital cortex.

**Figure 4/Table 2:**
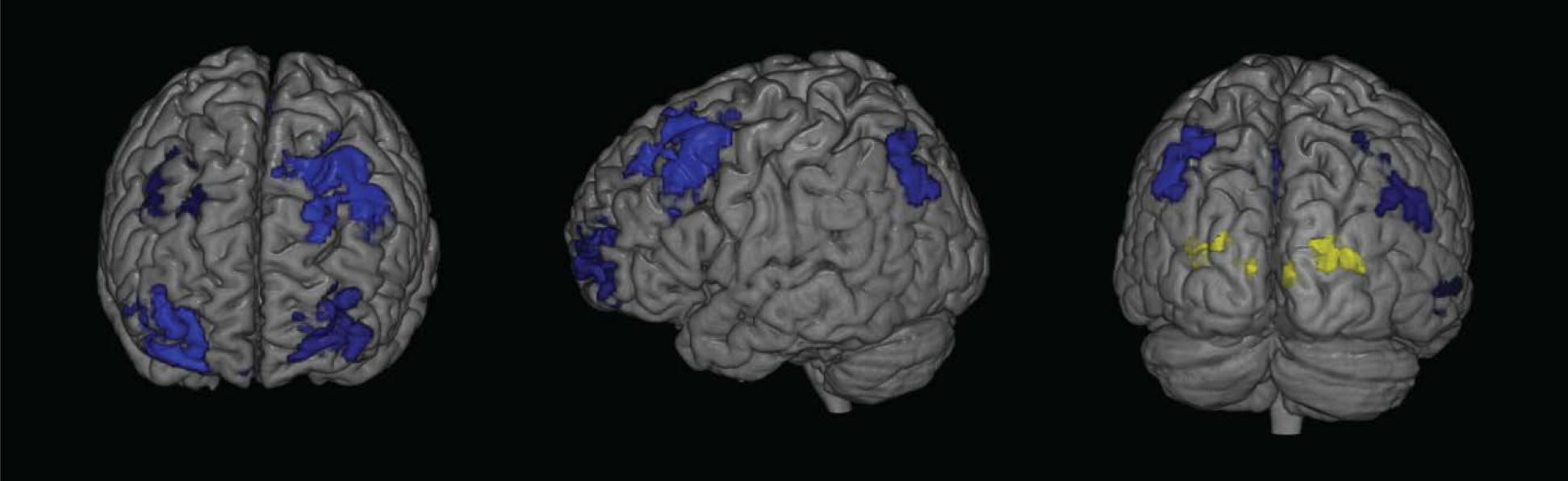
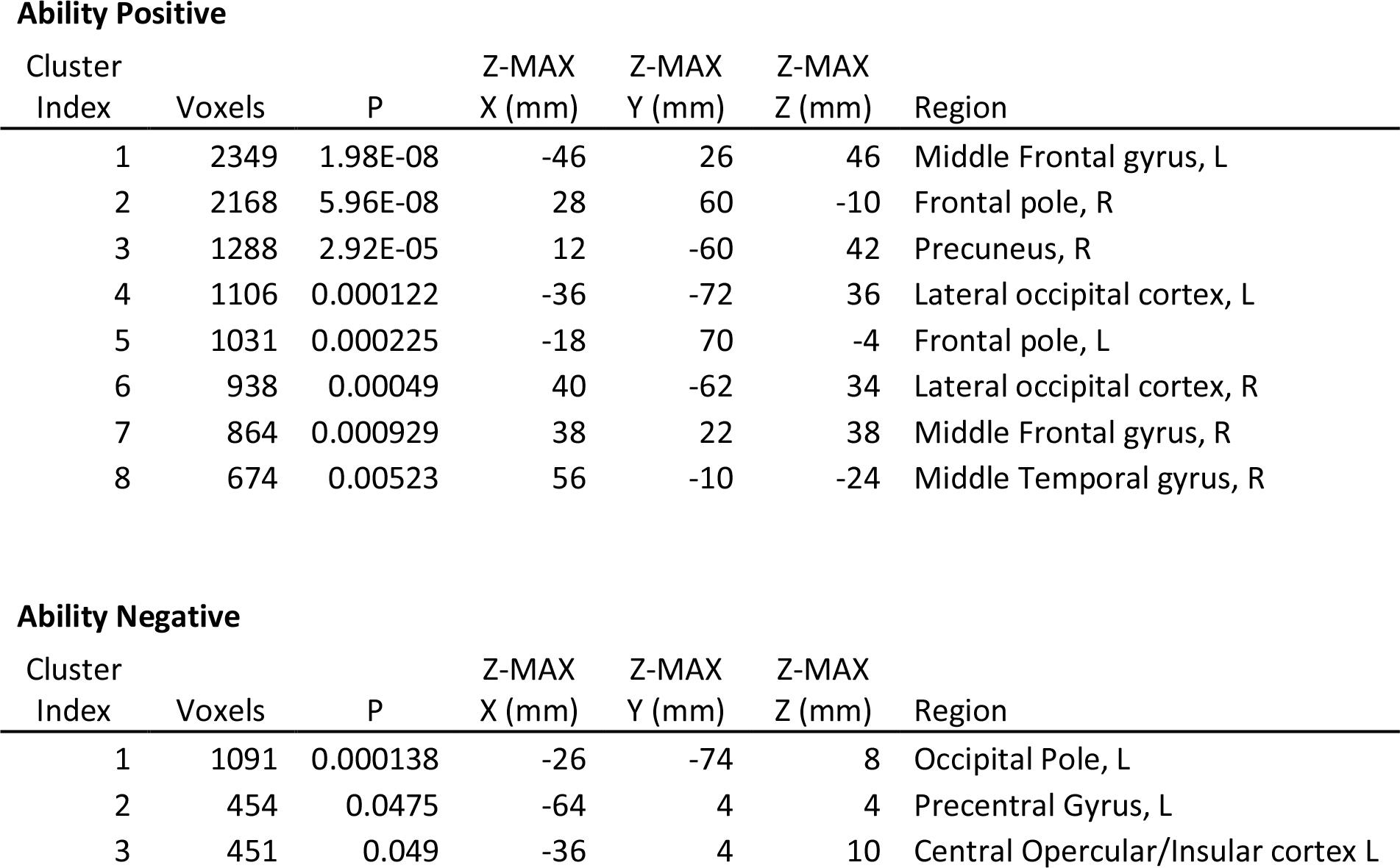
Spatial activity maps showing clusters of greater activity in individuals of higher fluid reasoning ability (blue) and clusters with significantly less activity in individuals with higher ability (yellow). Peak activations of each of the clusters are shown in the table.

### Conjunction and disjunction of intra-and inter-individual dimensions

Next, we formally examine the (dis)similarities between the inter-and intra-individual dimensions. This is of interest for both methodological and intrinsic reasons. First, the extent to which these dimensions *differ* will illustrate the methodological perils of conflating the inter-and intra-individual dimensions. To examine to what extent these two dimensions differed, we examine where in the brain the activity associated with differences in item difficulty was greater than the activity associated with differences in ability, and vice versa. The former contrast yielded no significant regions of interest, suggesting there are no regions of the brain that are more active for more difficult trials that aren’t also more active in people with higher ability. However, the converse contrast (where in the brain are individual differences in ability greater than differences due to item difficulty) showed widespread bilateral regions where the differences as a function of ability where greater than the differences associated with difficulty (Figure 5). These analyses suggest that neural patterns as a function of individual differences seem to be more spatially distributed than intra-individual differences as a function of task difficulty. This is the case despite the fact that the items ranged from extremely easy (made correctly by every person) to extremely hard (not being made correctly by any participant).

**Figure 5:**
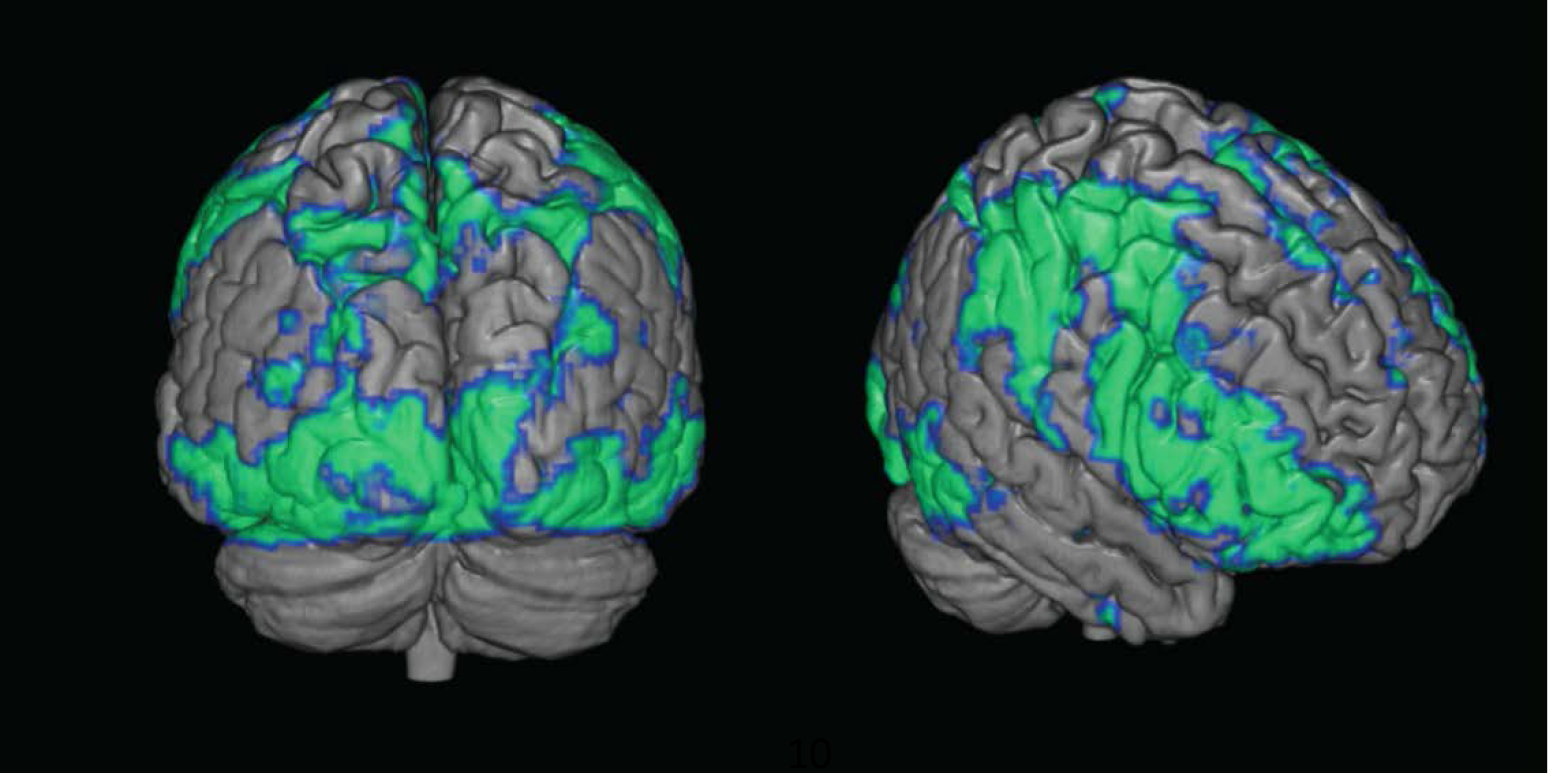
Regions showing greater activity as a function of individual differences in ability than as a function of difficulty. This contrast clearly illustrates the necessity of separating the two sources of variation.

The second question we can ask is not where these two dimensions differ, but where they overlap. It has long been known, although often ignored, that only in highly specific circumstances can we infer intra-individual processes from inter-individual differences (or vice versa). This inference is only valid when a process is *ergodic* (Molenaar, 2004). Ergodicity implies that the statistical characterization of within-subject variation is (asymptotically) identical to the variation at the level of the group (Molenaar, 2004; Molenaar and Campbell, 2009), which is very unlikely for most psychological constructs (Kievit et al., 2013). Although ergodicity is usually framed within the context of (natural) variation over time, it can be equally useful to describe intra-individual differences in task complexity (in which case it is closely related to *inter-individual measurement invariance,* e.g. see Adolf et al., 2014). Although some neuroimaging techniques have been developed that can test for ergodicity in neuroimaging for specific designs, e.g. in event-related connectivity (Gates et al., 2011), it is still a relatively neglected topic. Generally, these techniques examine *‘global* ‘ergodicity, that is, are the observed patterns as a whole identical for inter and intra-individual comparisons. We here propose a more lenient, but conceptually useful, form of ergodicity for neuroimaging: Where in the brain does intra-individual manipulation yield *the same differential activity* as that which characterizes inter-individual differences on that task (e.g. Sliwinski et al., 2010; Raz and Lindenberger, 2011; Voelkle et al., 2014)? We refer to this pattern as *local ergodicity.*

Such an analysis is useful for a variety of reasons. First, it forces us to make explicit the distinction between intra-and inter-individual differences, an issue often neglected in cognitive neuroscience. Second, and more importantly, regions that are relevant in explaining both dimensions are more likely to be fundamental to the cognitive phenomenon of interest. As the convergence of inter-and intra-individual this patterns is a non-trivial requirement, cases where it does apply they are likely to tell us something about the mechanics underlying the phenomenon of interest. Conversely, the lack of overlap between the two domains may be informative in terms of the cognitive insight gained into the phenomenon. If the neural systems underlying inter-and intra-individual differences are clearly separable, then explanatory accounts of the two dimensions should differ accordingly. Much like the relationship between speed and accuracy can be explained by qualitatively different mechanisms within people (a negative relationship, due to a speed accuracy trade-off) and between people (a positive relationship, due to the positive manifold), violations of ergodicity in neuroimaging may lead to more refined proposed neural mechanisms that capture each dimension separately.

**Figure 6:**
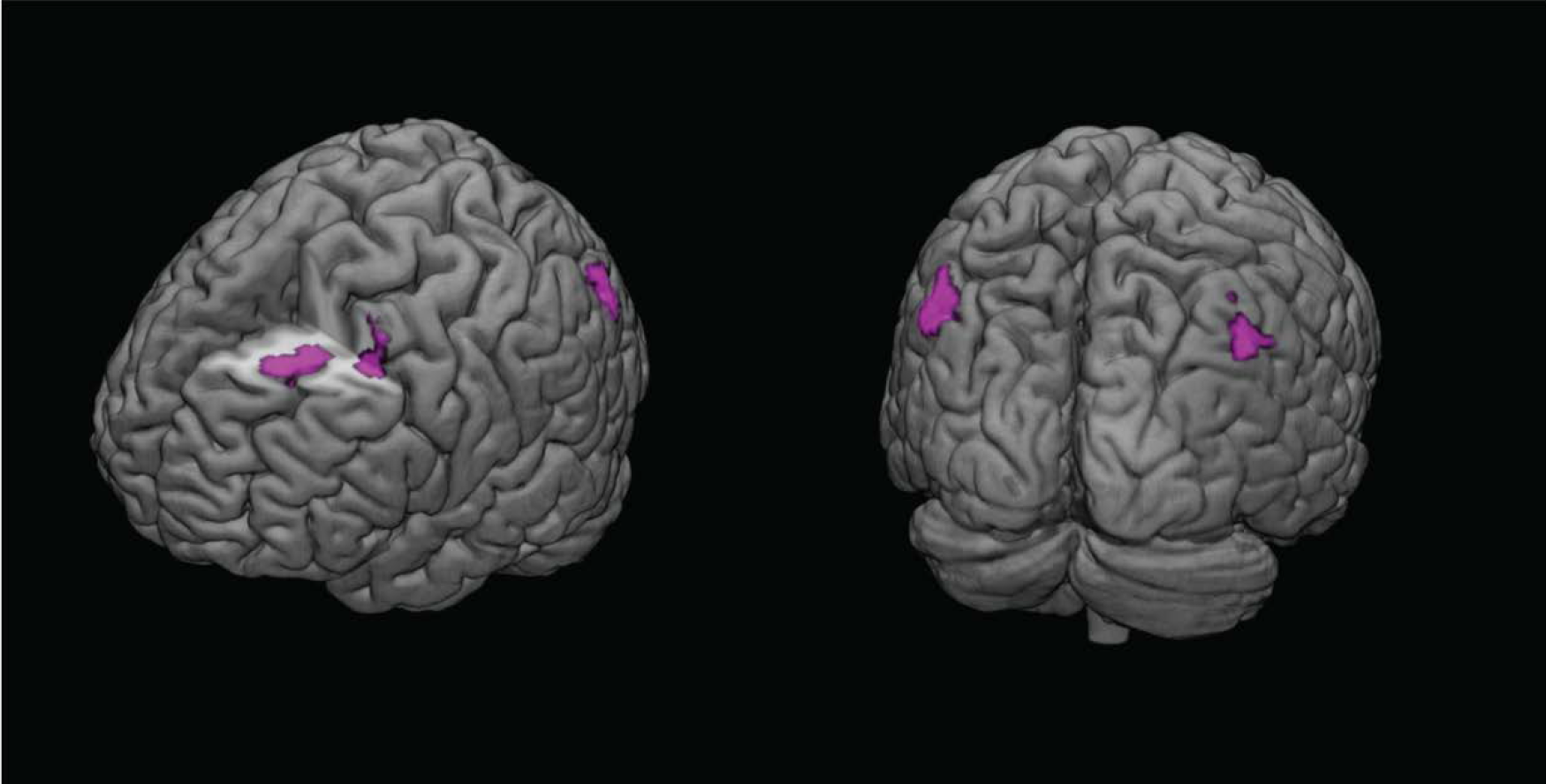
Locations that show evidence of neural ergodicity. These three clusters (bilateral angular gyri, bilateral precuneus and left superior frontal gyrus) showed increased activity both for more difficult items and for individuals of greater ability.

To test for neural ergodicity, we performed a conjunction analysis (Nichols et al., 2005) based on the statistical maps of the first two analyses (intra-individual differences in difficulty shown in Figure 3 and inter-individual differences in ability shown in Figure 4). Figure 6 shows the parietofrontal regions that are more active both as a function of increasing item difficulty and increased ability, and can be said to display *local neural ergodicity.* These regions can be described broadly as three clusters: The bilateral angular gyri in the superior parietal cortices, the bilateral precunei and the right middle and superior frontal gyri. Most notably the left lateral frontal region shows a striking convergence with a recent finding that greater connectivity of the LPFC predicts better fluid reasoning performance (Cole et al., 2012). In our final analysis we use IRT analysis to reexamine the ongoing debate on neural efficiency.

### Correcting for individual differences: Neural (in)efficiency

Finally, we examine how we may use the parameters estimated in our IRT model to get a better grasp of possible mechanisms differentiating individuals. Appropriately taking into account both intra-and inter-individual variation is important for a particular, hotly contested, hypothesis, namely that of *neural efficiency* (Haier et al., 1988; Neubauer et al., 2002; Neubauer and Fink, 2009; Poldrack, 2014). In its most common form, neural efficiency is the claim that individuals of higher ability show less activity during cognitive tasks because they ‘display a more focused cortical activation during cognitive performance resulting in lower total brain activation than in less intelligent individual’ (Neubauer et al., 2002, p. 515). However, this concept has been challenged recently for being little more than a tautological redescription of the data (cf. Poldrack, 2014, p. 2) such that ‘those of higher ability’ finding the same task ‘less hard work’. A more relevant question, we argue, is to compare individuals of different abilities when they are forced to ‘work equally hard’ (We note that Neubauer and Fink, 2009, do mention task complexity as a possible moderator of neural efficiency, e.g. pp. 1013). In other words, to meaningfully study differences in the processes that occur when individuals are being challenged cognitively (high difficulty), we must control for baseline differences in ability. Doing so, we can study the more relevant question of whether differences between individuals in fluid reasoning ability are associated with different cognitive patterns that would be suggestive of different cognitive strategies when performing items of equal difficulty. Item response theory allows for an easy way to separate difficulty and ability in precisely this manner.

For this analysis, we must ensure we select a subset of items that are equally difficult for all individuals. To do so we can simply subtract individual ability scores (theta) from the difficulty of each item (beta), to get a *corrected difficulty score* for each item, for each person. Next, we select a subset of all 72 items for each individual, such that that the range of items is equal in difficulty across all individuals. If after controlling for individual differences we find different patterns of activation, this would suggest that people who score more highly on Raven's matrices don’t simply make more items correctly: They recruit different neural networks than those with lower ability, even when making items of equal (subjective) difficulty.

We selected a subset of 30 items for every individual such that the *mean corrected difficulty* (defined as the difficulty of the items minus the ability of the participants) of those items was equal for every individual. We compared these subsets across every pair of individuals and found no significant differences in corrected item difficulties (all p's>0.069). However, significance tests are poorly equipped to quantify the absence of effects (Wagenmakers, 2007), so we reran this comparison using default Bayesian t-tests (Morey and Rouder, 2013). This analysis showed no evidence for significant differences and considerable evidence for an absence of such differences (mean BF_01_= 2.95, max BF_01_= .92). Next, we repeated the analysis shown in Figure 3 by first; calculating per subject, a dummy contrast for the trials that where equated for difficulty (e.g. equally difficult given the capacity of the subject) and using these in a FLAME analysis in which we entered ability estimates (theta) to answer the follow question: Which regions (if any) are more active in individuals with higher ability, and which regions (if any) are more active in individuals of low ability, given that we have equated the subjective difficulty of the items included in the analysis? Figure 7 shows both results: In cyan, we can see five main clusters that are more active in individuals who have greater fluid reasoning ability^1^. These clusters include the Anterior Cingulate Cortex, a small posterior cluster bordering on the right supramarginal gyrus and a most pronounced frontal set of clusters in the orbitofrontal cortex and the dorsolateral prefrontal cortex, extending to the left frontal pole. Moreover, it shows that, contrary theories of neural efficiency, in our sample individuals with greater ability generally show *greater* frontal and prefrontal activity when performing items of the same subjective difficulty. Second, we examine whether there are any regions that are more active in individuals with lower fluid reasoning ability. This analysis yielded a single, left-lateralized cluster, shown in yellow/red, bordering the extrastriate cortex in the superior occipital lobe, on the intersection between Brodmann area 18 and 19.

**Figure 7/Table 3:**
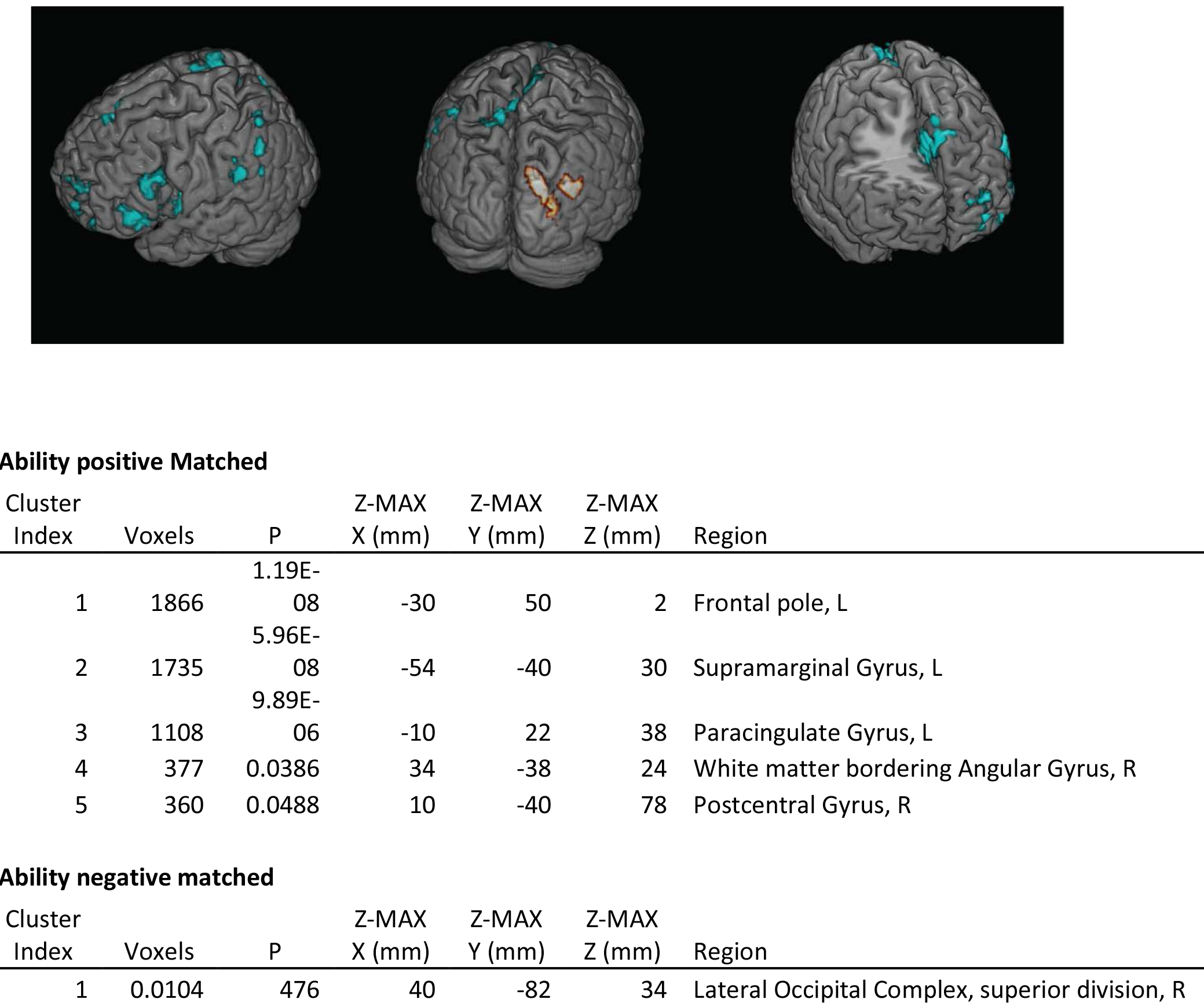
Greater activity in people with higher fluid intelligence (cyan) and lower fluid intelligence (yellow/orange) once item difficulty has been equated across individuals. This pattern suggests that, beyond the unidimensionality of the behavioural variables, there exist individual differences in neural responses during fluid reasoning items. Using Neurosynth suggests these differences are compatible with a more memory-and rule maintenance based strategy for individuals with high fluid intelligence compared to a more visual, object-oriented strategy by individuals with lower ability.

This analysis shows there are differences in neural activity between individuals of higher and lower fluid reasoning ability even when they perform equally challenging tasks. Next we may ask what these differences reveal about the cognitive processes underlying these differences. Inferring mental states and styles from neuroimaging patterns is notoriously complex (Henson, 2005; Poldrack, 2006). However, we can use Neurosynth (http://neurosynth.org/, Yarkoni et al., 2011), an automated metaanalysis tool using automated analysis of neuroimaging data and keyword frequency across more than 11,000 studies. This database makes it possible to provide a descriptive heuristic of cognitive states possibly associated with activity patterns, based on what has been reported in previous studies. Specifically, we can use the peak activations as shown above for both difficulty and ability to compute the posterior probability of a certain keyword being mentioned with high frequency (>1 in 1000) in any of the articles included in the database that also report activity in that cluster. Performing this analysis for the peak cluster above shows that the peak activation of the cluster that is more active in people with lower gf (X= 40mm, Y=-82mm, Z=34mm) has a posterior probability of .79 with the keyword ‘objects’. This is in line, tentatively, with the hypothesis that individuals with lower fluid intelligence, on average, rely on more purely visual strategies when performing fluid reasoning tasks. In contrast, the peak cluster for individuals with higher fluid intelligence (X= -30mm, Y=52mm, Z=2mm) is instead associated with keywords such as ‘memory’ (posterior probability=.7), ‘retrieval’ (posterior probability=.74) and ‘maintaining’ (posterior probability=.83). This is compatible with the hypothesis that individuals with higher fluid intelligence rely more on more frontal and prefrontal regions associated with memory and rule based strategies to solve fluid reasoning items. However, specific paradigms (e.g. selective disruption of via TMS of frontal versus occipital regions) would be necessary to support this hypothesis, as Neurosynth in isolation can only provide part of the picture (cf. Yarkoni, 2015a, 2015b). If our inference above is correct, higher gf individuals would be more adversely affected by lateral frontal stimulation whereas lower gf individuals would be more affected by high visual disruption.

Together, these results suggest that when individuals who vary in ability have been matched to perform tasks that are equally difficult, there are noticeable differences in neural patterns, such that people of higher ability show relatively more left lateralized prefrontal activity, whereas individuals with lower ability show more right lateralized higher visual activity. These findings do not support the general neural efficiency hypothesis, instead suggesting more complex *qualitative* differences between high and low ability individuals. Our analysis shows how we can refine this question using psychometric techniques such that neuroimaging can reveal individual differences beyond a well fitting unidimensional of purely behavioural data. We agree with Poldrack (2014) that neural efficiency as it is often operationalized will rarely be the question we are interested in, and suggest that psychometric techniques are more commonly used to refine the question at hand.

## Discussion

In this study we decompose two distinct but equally important dimensions of fluid intelligence: Intra-individual differences in neural responses to items of varying difficulty and inter-individual differences in neural activity for individuals of differing fluid reasoning ability. We use a parametric IRT model to show that greater ability in fluid intelligence is associated with broad, bilateral increases in activation of fronto-parietal regions, whereas increases in activity within individuals as a function of difficulty are associated with a more focal set of regions including the angular gyri and the precunei. In addition to these differences, we find a subset of three cortical systems, namely bilateral parietal, bilateral middle frontal and bilateral prefrontal gyri, which show increases *both* as a function of difficulty and as a function of increased ability. We propose to describe this convergence of intra-and inter individual neural responses as *neural ergodicity.* The importance of the question of ergodicity for neuroscience is increasingly being realized, as recent work suggests that even within a relatively well-controlled network analysis, ergodicity is violated (i.e. differences in network connectivity within and between subjects do not converge, Medaglia et al., 2011). Moreover, this subset of regions may reflect key processes emphasised by cognitive theories that link inter-and intra-individual processes in intelligence (e.g. Van Der Maas et al., 2006; Kovacs and Conway, 2016).

Crucially, we show that IRT can be used to equate item difficulty across a sample of individuals that differ in ability, thus making the neural comparisons more psychologically meaningful and comparable. Doing so suggests that high ability individuals show more frontal activity whereas individuals of lower ability show more occipital activity. These findings are compatible with, although certainly not conclusive of, strategy differences in reasoning style for individuals of higher and lower ability. Crucially, our approach can be extended to related cognitive domains that study both ability and difficulty levels, such as working memory (e.g. Pessoa et al., 2002 versus (Osaka et al., 2003) cognitive control (e.g. Botvinick et al., 2001 and Hester et al., 2004) and many more.

Applying IRT in this context provides not only a cautionary note in conflating the two dimensions, but illustrates how neuroimaging data may allow us to better understand the mechanisms of individual differences: Although the fit of the Rasch model to the behavioural responses was good (suggesting unidimensionality of the behavioural responses), a more detailed investigation of the neural responses showed residual differences between individuals of high and low ability informative of the underlying cognitive processes. Practically, the dissociation of intra-and inter-individual dimensions suggest that many study designs may not be tailored to answer the question of interest in the most efficient, or even most accurate, way - Findings of individual differences will depend, in part, on the range of item difficulties presented, and findings of parametric difficulty will depend on the mean ability and range of the population being studied.

Although the current approach represents a step forward in modelling the con-and divergence of two psychologically relevant dimensions, we are aware that we implicitly assume homogeneity in several other potentially dimensions. For instance, our study focuses on an age range (18-30) within which fluid intelligence is relatively stable. This means that in a sample with a larger age range, there is possibility that the neural systems underlying individual differences in fluid intelligence will be distinct from individual differences (of the same magnitude) seen in our sample. Future research may extend these findings by the better integration of the temporal dynamics of the cognitive processes underlying fluid reasoning. Recently developed psychometric models have combined intra-individual processes as an information-accumulation process with inter-individual differences in ability (van der Maas et al., 2011). Such models could be fruitfully combined with similar integrative developments in neuroimaging methods that combine the high spatial resolution of fMRI with more time-sensitive methods, leading to (combined) M/EEG or fMRI/EEG.

The central role of fluid reasoning in human cognition means that the potential payoffs of a better (mechanistic) understanding are considerable. A better mechanistic understanding of fluid reasoning is essential for the promise of targeted behavioural or neurological training delaying deterioration of fluid reasoning during old age (e.g. Salthouse, 2009) or recovery of fluid reasoning associated difficulties after strokes or lesions (Duncan et al., 1995; Woolgar et al., 2010; Barbey et al., 2014) and the promise of clinical assessment of cognitive faculties by means of standardized neuroimaging tests (Allen and Fong, 2008). However, before we can get close to a mechanistic level of understanding require to achieve these ambitious goals, careful study of two fundamentally different dimensions of psychological variation is crucial: The inter-individual domain and the intra-individual domain. Over 50 years ago, Cronbach (1957) referred to these domains as ‘the two disciplines of scientific psychology’ and it has been questioned to which extent these two domains have been brought closer together since (Borsboom et al., 2009; Voelkle et al., 2014). By using formal models to best capture inter-and intra-individual phenomena in neuroimaging studies, the two sub-disciplines of cognitive neuroscience may more quickly, and more fruitfully, converge.

### Materials and Methiods

*Participants:* 37 participants (19 female) with normal or corrected-to-normal vision participated. The participants were tested in accordance with the ethical guidelines of the American Psychological Association, and the study was approved by the University of Amsterdam Ethics Committee, and received a financial reward for their participation. Prior to analysis, three subjects were excluded because of excessive motion (N=1) or scanner malfunction (N=2), leaving 34 subjects (17 female). The final sample ranged in age from 18 to 30 (M=23.4, SD=2.8).

In the scanner, subjects performed a total of 72 Raven's matrix items, drawn from the Standard Progressive Matrices (36 items from sets C, D and E, Raven 1960) and Raven's Advanced Progressive Matrices items (Raven, Court, & Raven, 1996). Figure 1 shows a stylized example of an easy item and a difficult item. The eight-option items were adapted for use in the scanner. Ravens matrices are considered good measurements of fluid reasoning ability and figure centrally in psychometric analyses of general intelligence (Carpenter et al., 1990). The experiment was programmed using Presentation^®^ (Neurobs, 2011). Participants viewed the screen (61 cm x 36 cm) on which the stimuli were presented via a mirror mounted on the head coil. Participants had a four-button box in each hand to respond to the eight clearly marked answer options. Prior to the first scan subjects were able to practice pressing the buttons with visual feedback to ensure correct response mapping.

All raw behavioural data and an analysis script written in R (Team, 2014) are freely available on Figshare (both behavioural data and basic demographics https://figshare.com/articles/Rawdatafiles/2077552 as well as the R code https://figshare.com/articles/Analysis_code_written_in_R/2077555). The neuroimaging first level data and key contrasts are freely available on Neurovault (Gorgolewski et al., 2015) at http://neurovault.org/collections/1160/.

### Procedure

Prior to the scanning session, subjects read instructions and performed 12 practice trials (not used in the study) to ensure they understood the task. After ensuring the instructions were clear, participants were placed in the scanner. Each block consisted of 12 Raven's matrices, interspersed by a 16 second inter-trial interval, with a maximum 30 second response window for each item. The blocks were pseudo-randomized such that each of the six blocks contained 12 fixed items spanning the complete range of difficulty (from easy to difficult), but were randomized within each block. This ensured that subjects did not ‘give up’ because trials became increasingly complex within or across blocks.

### Image acquisition and pre-processing

Imaging data were obtained at the University of Amsterdam Spinoza Centre for Functional Magnetic Resonance Imaging using a 3-T Philips Achieva TX scanner using an 8-channel head coil. During the presentation of the Raven tasks we recorded BOLD-MRI (GE-EPI, TR=2346 ms, TE=30 ms, FA=90°, transversal recording, FOV=200^2^ mm, matrix size=80^2^, 39 slices, slice thickness=3, slice gap=0.3, ascending acquisition). We also acquired a high-resolution anatomical recording (3DT1, TR=8.1 ms, TE=3.74, FA=8°, FOV=240*220*188 mm, voxel size=1 mm^3^) for normalization purposes. Foldable foam pads were used to minimize head motion. Data were analysed using FSL (FMRIB's Software Library, www.fmrib.ox.ac.uk/fsl, MATLAB (Version 7.10.0, The Mathworks, Inc., Natick, MA, USA), and R (Team, 2014). Functional data were analysed using FEAT (FMRI Expert Analysis Tool Version 5.98), in which we performed motion correction, slice time correction, spatial smoothing (5mm) and low pass filtering (100 s). We generated explanatory variables for each individual presented item of the Raven's progressive matrices using the double gamma model of the hemodynamic response function. This yielded 12 explanatory variables (EV's) per run. These EVs were subsequently combined using a model in which we specified both the mean activation level and the item difficulty for each item. This yielded an estimate, per subject, of the extent to which activity of voxels differed across items varying in item difficulty. At the between-subject level we specified a model in which the average activity of the covariate fit from the fixed effects pooling stage was entered and in which the ability of the individual subjects was included as a predictor, so we could the estimate the effects of item difficulty independently of subject ability. Higher-level analysis was carried out using FLAME (FMRIB's Local Analysis of Mixed Effects) stage 1 and stage 2 with automatic outlier detection (Woolrich et al., 2004; Woolrich, 2008). Statistics were thresholded using cluster-based correction at z=2.3 and a corrected cluster significance threshold of 0.05 (Worsley, 2001)

## Footnotes

1 Note that due to a smaller subset of available Items we did not model the Intercept for the dummy variables at subject levels, so the t-values at the individual level are higher than the t-values in the analysis presented in Figure 2.

## Acknowledgements

RAK, HSS, LW and DB conceived of the experiment. RAK programmed the experiment, performed the behavioural analysis, main authored the article and created the figures. HSS and RAK performed the neuroimaging analysis. All authors contributed to the final manuscript.

